# Cell-specific delivery of GJB2 restores auditory function in mouse models of DFNB1 deafness and mediates appropriate expression in NHP cochlea

**DOI:** 10.1101/2024.12.24.630240

**Authors:** Maryna V. Ivanchenko, Kevin T.A. Booth, K. Domenica Karavitaki, Larisa M. Antonellis, M. Aurel Nagy, Cole W. Peters, Spencer Price, Yaqiao Li, Anton Lytvyn, Andrew Ward, Eric C. Griffith, Sinisa Hrvatin, Michael E. Greenberg, David P. Corey

## Abstract

Mutations in the *GJB2* gene cause the most common form of human hereditary hearing loss, known as DFNB1. *GJB2* is expressed in two cell groups of the cochlea—epithelial cells of the organ of Corti and fibrocytes of the inner sulcus and lateral wall—but not by sensory hair cells or neurons. Attempts to treat mouse models of DFNB1 with AAV vectors mediating nonspecific *Gjb2* expression have not substantially restored function, perhaps because inappropriate expression in hair cells and neurons could compromise their electrical activity. Here, we used genomic chromatin accessibility profiling to identify candidate gene regulatory elements (GREs) that could drive cell-type-specific expression of *Gjb2* in the cochlea. HA-tagged GJB2, delivered to a conditional knockout model in an AAV vector with GRE control of expression, was localized to the appropriate cell types, prevented the cochlear degeneration observed in untreated knockout mice, and partially rescued hearing sensitivity. In a *Gjb2* partial knockdown mouse model, such exogenous GJB2 prevented degeneration and completely restored hearing sensitivity. We tested control of expression by these GREs in nonhuman primate cochleas and found that vector-delivered human GJB2.HA was located in the appropriate cell types and caused little or no reduction in hearing sensitivity. Together, these findings suggest that GRE-mediated expression of *GJB2* could prevent hearing loss in DFNB1 patients.

## Introduction

Hearing loss is the most common sensory loss ^1,2^, affecting 2-3 in 1000 children born in the United States. Although more than 180 genes have been causally linked to hearing loss^3^, a single form of recessive, non-syndromic hearing loss, DFNB1, is responsible for up to 50% of congenital deafness worldwide outside of sub-Saharan Africa^4,5^. In the United States alone, about 3,500 children are born each year with mutations in both alleles of the causative gene, *GJB2*^4–7^. Many are born with profound hearing loss, but two-thirds have some residual hearing at birth and the majority of those lose hearing over the next few years^8^, suggesting that a developmental window exists for therapeutic intervention.

*GJB2* encodes the gap junction protein GJB2/connexin26. In the cochlea, *GJB2* is expressed in two cell groups: an epithelial system comprising supporting cells of the organ of Corti, epithelial cells of the inner and outer sulcus, interdental cells; and a connective tissue system comprising fibrocytes of the lateral wall and suprastrial zone, and basal cells of the stria vascularis^9–11^. GJB2 is absent from the sensory hair cells and the postsynaptic spiral ganglion neurons.

There are several hypotheses for why GJB2 is critical for the cochlea. In one, the K^+^ that enters hair cells through transduction channels and leaves through basal K^+^ channels is shuttled away from the organ of Corti by the epithelial system^9^. Failure to remove K^+^ from the extracellular medium would tend to depolarize hair cells and allow toxic Ca^2+^ influx through voltage-gated calcium channels. Another is that GJB2 in fibrocytes and strial cells is important for transporting K^+^ into endolymph and producing the endocochlear potential^9^. A third hypothesis suggests a required developmental role, perhaps involving glucose transport, as mice lacking GJB2 in the inner ear have profound loss of hair cells and supporting cells by P30^12–15^. If *Gjb2* is deleted after P6, the phenotype is much milder^16^; however, there remains a long-term requirement for GJB2, as hair-cell loss occurs after months even with deletion as late as P14^17^. A fourth hypothesis involves ATP- and connexin-dependent intracellular Ca^2+^ signaling in development^18^. Regardless of etiology, these studies point to a critical role for GJB2 in the early postnatal period in mice, with a sustained but less critical long-term function.

Adeno-associated virus (AAV) mediated gene addition to neonatal *Gjb2* conditional knockout mice might be expected to rescue this pathology. Yet this approach has thus far failed to substantially rescue function^12,19,20^. There may be various reasons for this failure—whether the capsid transduces all the appropriate cells, whether regulatory elements prevent expression in inappropriate cells, whether the vector dose is adequate, or whether the vector is delivered soon enough to prevent cochlear degeneration—but each study has used different combinations of these elements and their relative importance for functional rescue has not been systematically studied.

Here we describe a successful gene replacement approach for DFNB1-related hearing loss employing ATAC-seq (Assay for Transposase-Accessible Chromatin using sequencing) identification of GREs to limit off-target expression of GJB2 and fully restore hearing thresholds in a *Gjb2* mutant mouse model. Furthermore, we demonstrate that our vector effectively targets GJB2-expressing cells in a non-human primate (NHP) model without inducing toxicity. This study is the first to achieve significant rescue in a *Gjb2* mouse model of DFNB1 deafness while also providing safety and tolerability data in an NHP model.

## Results

### GJB2 expression under a ubiquitous promoter can be lethal

We previously found that several capsids, including AAV9-PHP.B and AAV-S, efficiently transduce the cells in neonatal mouse cochlea that natively express *Gjb2*, based on expression of a GFP marker gene driven by a ubiquitous promoter^21–23^. To determine whether we can also express *Gjb2* in those cells, we packaged the coding sequence for mouse GJB2 (mmGJB2) in the AAV9-PHP.B vector under the control of the CBA promoter (AAV-CBA-mmGJB2; **Fig. 1a**) and injected the vector into the cochleas of neonatal (P0 or P1) mice through the round window membrane (RWM). Ubiquitous expression of mmGJB2 was often lethal. Mice injected with AAV-CBA-mmGJB2 exhibited severe shaking and almost all animals died by P5 (**Fig. 1b).** To determine whether the lethality was a consequence of GJB2 activity, we constructed a similar vector in which the GJB2 sequence contained the null mutation 35delG, which leads to a frameshift and stop (AAV-CBA-mmGJB2null). Mice injected with this vector survived until at least P30, the last time point evaluated (**Fig. 1b**), suggesting that lethality is a consequence of aberrant gap junction coupling.

**Figure 1.**
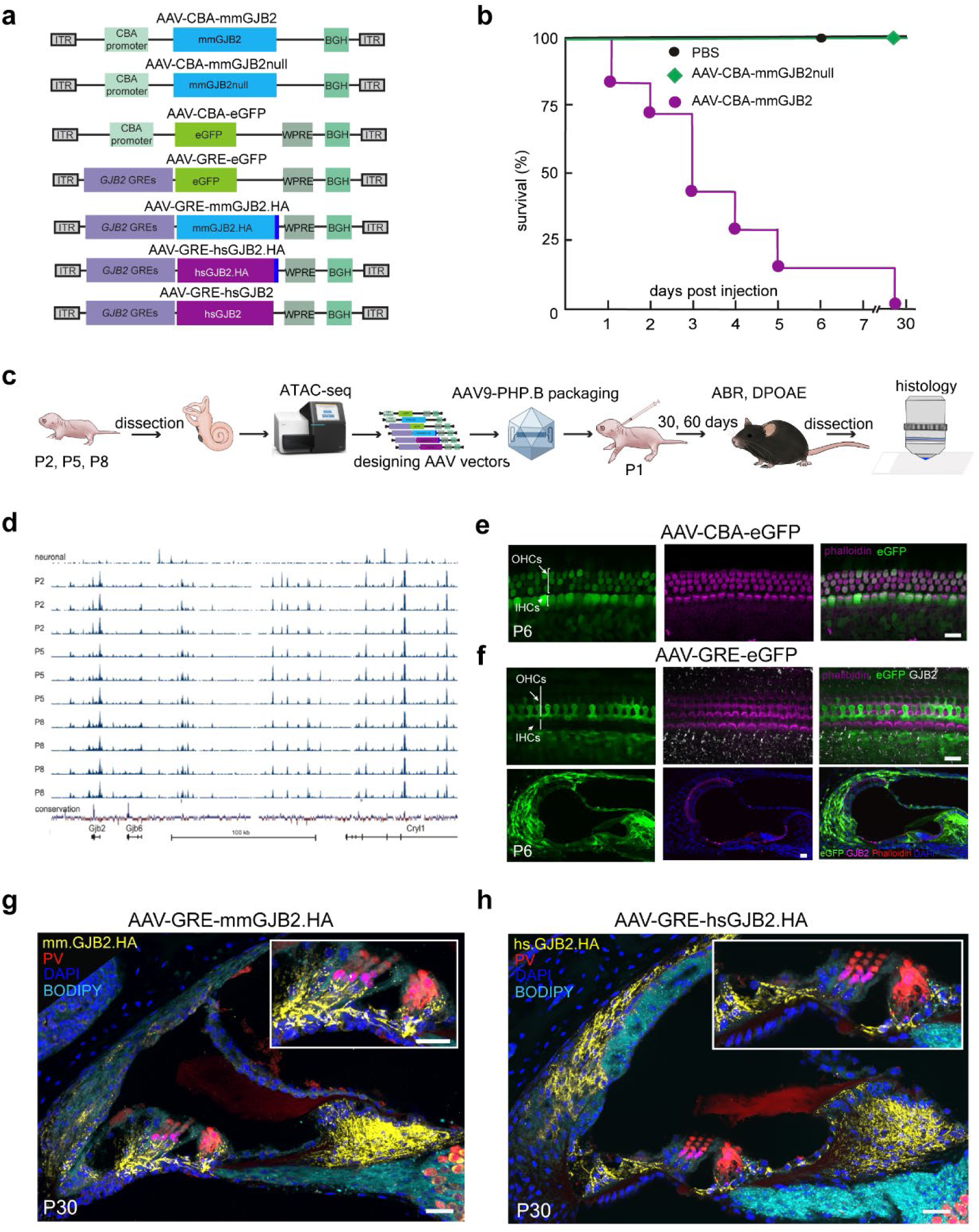
Gene regulatory elements restrict viral GJB2 expression. **a.** Schematics of the AAV vectors used in the study. All vectors are single stranded. ITR: inverted terminal repeat, CBA: hybrid CMV enhancer/chicken β-actin promoter, HA: hemagglutinin tag, WPRE: woodchuck hepatitis virus post-transcriptional regulatory element, BGH: bovine growth hormone poly A signal. **b.** Survival after vector injection at P1. Mice injected through the round window membrane with PBS survived to at least P6, the last day evaluated. Of seven mice injected with AAV-CBA-mmGJB2, six died by P5. Mice injected with vector encoding a null mutant GJB2 (35delG) lived to at least P30. **c.** Overview of the experimental design. **d.** Identification of potential GREs with ATAC-seq. Ten cochleas from neonatal mice (P2-P8) were harvested and processed for ATAC-seq. Peaks representing open chromatin are shown for a ∼300 kb stretch of mouse genome near the *Gjb2* locus. Peaks for a neuronal sample are shown at the top; sequence conservation among mammalian genomes is plotted at the bottom. **e.** Expression of a reporter driven by a ubiquitous promoter. An AAV9-PHP.B vector (AAV-CBA-eGFP) expressing eGFP (green) under a CBA promoter efficiently transduced inner hair cells (IHCs), outer hair cells (OHCs) (magenta), supporting cells, and other cells in mouse cochlea. IHCs and OHCs were identified by fluorescent phalloidin (magenta). **f.** Cells transduced with an AAV vector expressing the eGFP marker gene under the control of GREs (AAV-GRE-eGFP). Notably, no eGFP was observed in hair cells when expression was controlled by the GREs. **g, h.** Cells transduced with an AAV vectors express*ing* mmGJB2.HA (**g**) or hsGJB2.HA (**h**) under the control of regulatory elements. Scale bars: 10 μm (**e, f**), 30 µm (**g, h**).

AAV vectors injected into the mouse cochlea at early postnatal ages can spread to the brain^24–27^. We reasoned that nonspecific expression of *GJB2* in neurons driven by the CBA promoter could cause neurons to become electrically coupled to each other and to neighboring glial cells. The high conductance could short-circuit neuronal post-synaptic potentials; if expressed in inhibitory neurons, GJB2 could increase brain excitability and cause lethality. Indeed, the AAV9-PHP.B capsid was selected for its ability to transduce brain cells in neonatal mouse^28^. Similarly, in the cochlea, nonspecific expression of GJB2 in hair cells could electrically couple them to neighboring supporting cells and attenuate receptor potentials, abolishing the auditory brainstem response (ABR). When we used the promiscuous CBA promoter to drive expression of eGFP (AAV-CBA-eGFP; **Fig. 1a**) and injected it into the inner ears of P1 mice, we observed high expression of eGFP by P6 in cells of the sensory epithelium, including the inner and outer hair cells (**Fig. 1e**; **Table 1**).

**Table 1.**
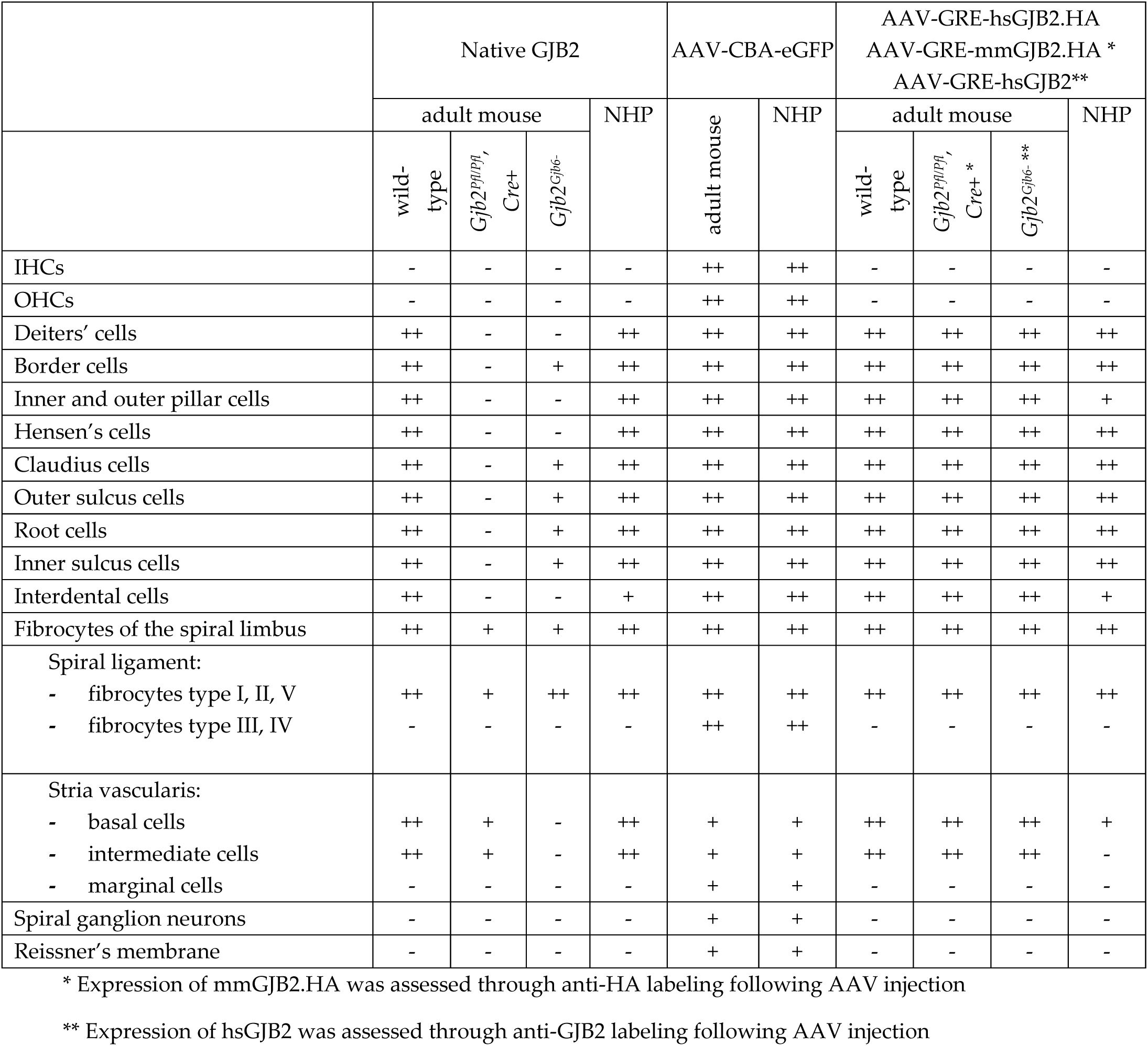
Expression of native GJB2, AAV-CBA-GFP, AAV-GRE-hsGJB2.HA and AAV-GRE-hsGJB2 in mice and NHPs.

### Gene regulatory elements restrict viral GJB2 expression

We therefore sought to limit virus-mediated expression to those cells that normally express *Gjb2*, using specific GREs. To identify GREs that control *Gjb2* expression in the mouse cochlea, we dissected whole cochleas from neonatal mice (age P2, P5 and P8), purified dissociated nuclei, and employed the Assay for Transposase-Accessible Chromatin with Sequencing (ATAC-seq)^29,30^ to reveal regions of open chromatin (**Fig. 1c**), as previously described^31,32^. ATAC-seq revealed several dozen such regions, both proximal and distal to the *Gjb2* gene (**Fig. 1d**). To restrict expression to the cochlear fibrocytes and epithelial cells that express *Gjb2*, we avoided regions that also appeared in a neuronal sample; to select regions more likely to control gene expression, we focused on those conserved among mammalian genomes (**Fig. 1d**). For more efficient translation to clinical use, we chose human sequences matching those of the identified mouse candidates.

We then constructed AAV vectors that encoded eGFP driven by our selected GREs (**Fig. 1a**) and injected them into neonatal mouse cochleas. We found that this vector (AAV-GRE-eGFP; **Fig. 1a**) resulted in a restricted pattern of eGFP expression that more closely matched endogenous expression of *Gjb2* (**Fig. 1f**; **Fig. 2a, c**). Notably, there was not expression in hair cells (**Fig. 1f**). Next, we injected neonatal cochleas at P1 with vectors encoding either mmGJB2.HA or hsGJB2.HA, adding a hemagglutinin (HA) epitope tag to the C-terminus to track virally expressed GJB2. We analyzed expression in cochleas at P30 using anti-HA staining (**Fig. 1g, h**). Consistent with normal GJB2 localization, HA-tag expression was detected predominantly in the spiral limbus, inner and outer sulcus epithelia, supporting cells, basal and intermediate cells of the stria vascularis, and fibrocytes of the spiral ligament (**Fig. 2c**). *GJB2* was not expressed in inner hair cells (IHCs) or outer hair cells (OHCs). Immunolabeling appeared concentrated into disc-like puncta at cell membranes, indicating that GJB2 was expressed in the appropriate cells and properly trafficked to the cell membrane to form gap junctions (**Fig. 1g, h**, **Table 1**). A similar expression pattern was observed for both mmGJB2.HA and hsGJB2.HA.

**Figure 2.**
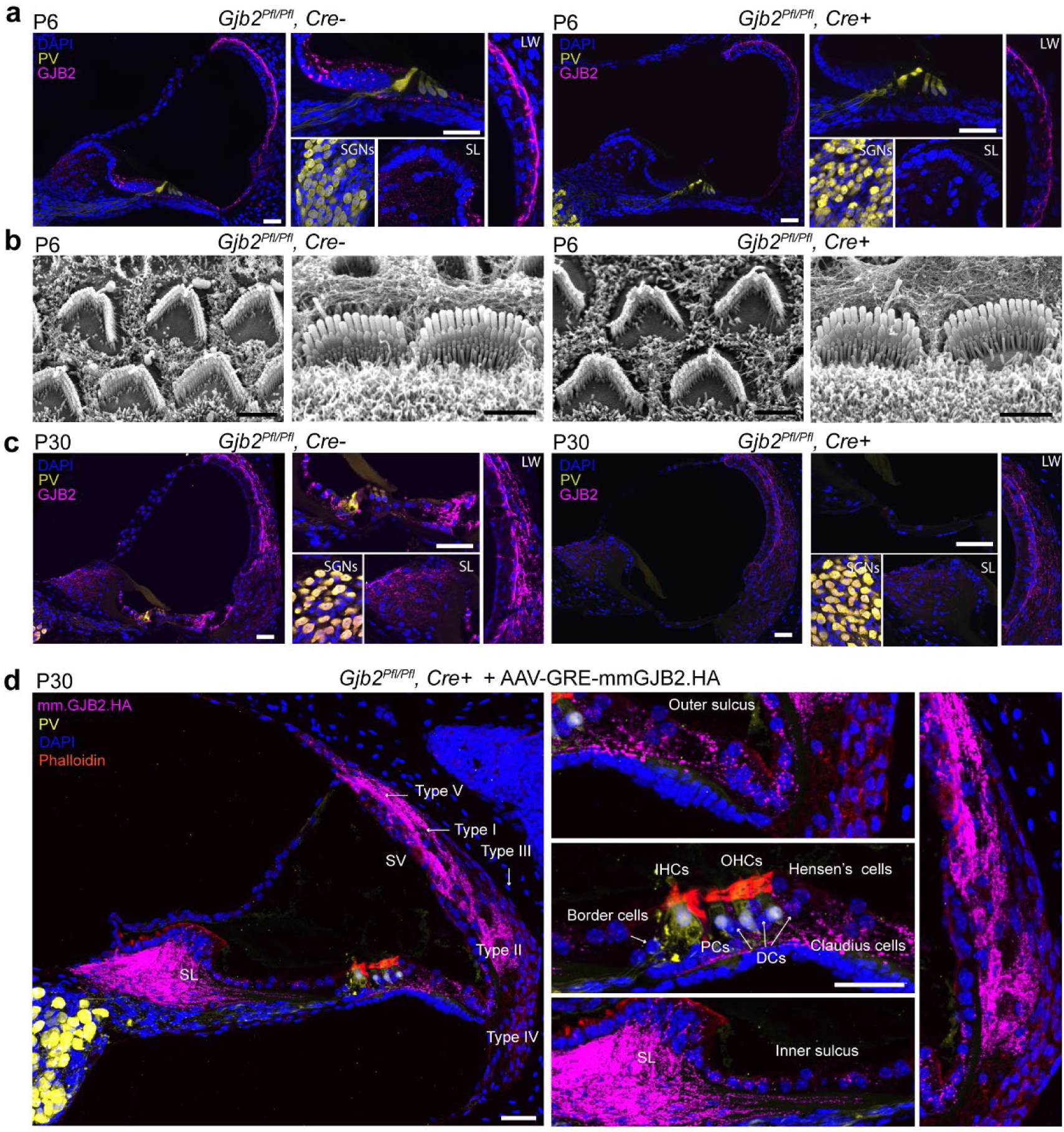
*Gjb2^Pfl/Pfl^, Cre+* conditional knockout mice at P30 exhibit degeneration of the sensory epithelium, and AAV.B-GRE-mmGJB2.HA delivery preserves cochlear morphology. a. Distribution of *GJB2* (*magenta*) in *Gjb2^Pfl/Pfl^, Cre-* control mouse cochlea (*left*) and *Gjb2^Pfl/Pfl^, Cre+* cochlea *(right)* at P6. In *Gjb2^Pfl/Pfl^, Cre- mice,* GJB2 was observed in Deiters’ cells, Hensen’s cells, Claudius cells, pillar cells, border cells, the epithelium of the inner and outer sulcus and the root cells of spiral prominence; in fibrocytes in the lateral wall and spiral limbus; and in basal and intermediate cells in the stria vascularis. Hair cells, labeled with an antibody to parvalbumin (PV; yellow) did not express *Gjb2* in control cochleas. In *Gjb2^Pfl/Pfl^, Cre+* conditional knockout mice at P6, anti-GJB2 labeling detected no signal hair cells or epithelial cells in the organ of Corti. **b.** At P6, scanning electron microscopy of hair cells showed normal bundle morphology in both *Gjb2^Pfl/Pfl^, Cre+* conditional knockout mice and *Gjb2^Pfl/Pfl^, Cre-* controls. **c.** Distribution at P30 of GJB2 in *Gjb2^Pfl/Pfl^, Cre-* (*left*) and *Gjb2^Pfl/Pfl^, Cre+ (right)* mouse cochleas. Conditional knockout mice had developed degeneration of the organ of Corti by P30. **d.** Exogenous (vector-delivered) mmGJB2.HA expression in P30 *Gjb2^Pfl/Pfl^, Cre+* conditional knockout mouse cochlea. Hair cells did not express mmGJB2.HA. It appeared in the epithelium of the inner and outer sulcus and the root cells of the spiral prominence, fibrocytes type I, II and V of the lateral wall and fibrocytes of the spiral limbus and strial cells, and these cells survived to at least P30. Hair cells, missing at P30 in untreated *Gjb2^Pfl/Pfl^, Cre+* mice, were present in the vector-treated cochlea. LW - lateral wall, SL - spiral limbus, SGNs - spiral ganglion neurons. Scale bars: 30 μm (**a, c, d**), 2μm (**b**).

### Hair cells and supporting cells degenerate in conditional knockout mice, resulting in profound deafness

To understand the phenotype of mice lacking GJB2 and to evaluate gene therapies in a model of DFNB1, we used mouse lines with little or no expression of the *Gjb2* gene. While humans with two null alleles of *GJB2* are viable, mice with constitutive deletion of both alleles are embryonic lethal^33^. To restrict deletion of *Gjb2* to a limited number of tissues, including cochlea, we first used a conditional knockout mouse model that deleted the entire coding sequence (*Gjb2^tm1Ugds^*)^34^; referred to here as *Gjb2^Pfl^*, and crossed it to *Sox10-Cre* mice^12,35^; referred to here as *Cre+*. Mice in which Cre recombinase deleted *Gjb2* (*Gjb2^Pfl/Pfl^, Cre+*) were compared to normal controls without Cre (*Gjb2^Pfl/Pfl^, Cre-*) (**Fig. 2a-c**).

We first characterized normal *Gjb2* expression in the cochleas of control mice by immunolabeling cryosections from neonatal (P6) and adult (P30) mice with antibodies against GJB2 and markers for hair cells and spiral ganglion neurons (SGNs) (**Fig. 2**). Consistent with previous reports^12,35^, cochleas at age P6 from *Gjb2^Pfl/Pfl^, Cre-* control mice showed prominent *Gjb2* expression in the inner and outer sulcus epithelia, supporting cells, the basal and intermediate cells of the stria vascularis, and fibrocytes of spiral ligament and spiral limbus (**Fig. 1a**). GJB2 was not detected in IHCs or OHCs. By P30, expression levels had increased in these regions (**Fig. 1c**, **Table 1)** aligning with earlier findings^12,36^.

We then evaluated expression in *Gjb2^Pfl/Pfl^, Cre+* knockout mice. At P6, there was no anti-GJB2 labeling in the organ of Corti, or more generally in the epithelial gap junction network that extends from the spiral limbus through the organ of Corti to the spiral prominence (**Fig. 2a**), validating the GJB2 antibody and confirming Cre- mediated knockout of GJB2 in these cells. Some anti-GJB2 signal was nevertheless present in the fibrocytes of the lateral wall and spiral limbus, suggesting that these cells do not strongly express *Sox10* and do not efficiently delete *Gjb2*, allowing some *Gjb2* expression. At P6, despite the absence of a GJB2 network in the organ of Corti, anti-parvalbumin staining showed normal hair cell morphology (**Fig. 2a**). Furthermore, no ultrastructural abnormalities in IHC and OHC bundle morphology were detected with scanning electron microscopy at P6 (**Fig. 2b**). By P30, however, *Gjb2^Pfl/Pfl^, Cre+* mice had developed degeneration of both hair cells and supporting cells of the organ of Corti (**Fig. 2c**).

We measured auditory thresholds by ABR and distortion product otoacoustic emissions (DPOAE) recording in both the *Gjb2^Pfl/Pfl^, Cre+* mutants and *Gjb2^Pfl/Pfl^, Cre-* control littermates at P30. The *Gjb2^Pfl/Pfl^, Cre+* mice showed profound deafness across all frequencies tested. *Gjb2^Pfl/Pfl^, Cre-* animals showed thresholds consistent with wild-type hearing (**Fig. 3**).

**Figure 3.**
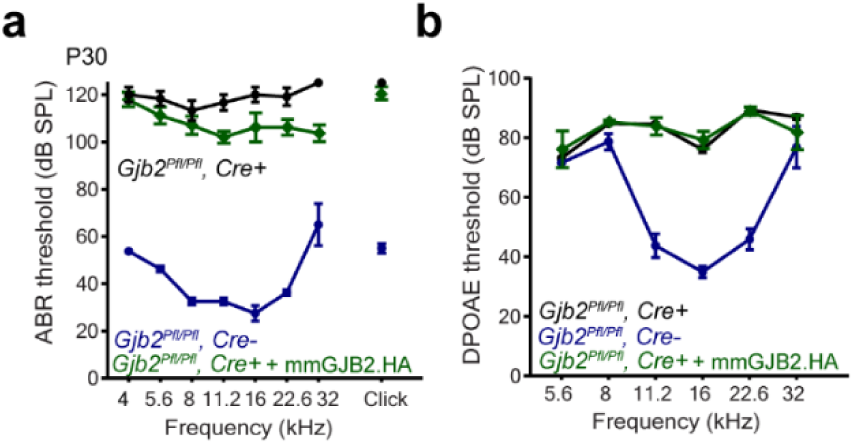
Deafness in *Gjb2^Pfl/Pfl^, Cre+* DFNB1 mouse models and rescue with vectors expressing GJB2.HA under control of GREs. **a.** Average tone ABR thresholds as a function of frequency and click thresholds for P30 uninjected *Gjb2^Pfl/Pfl^, Cre-* control mice (blue), uninjected *Gjb2^Pfl/Pfl^, Cre+* conditional knockout mice (black), and conditional knockout mice injected with AAV-GRE-mmGjb2.HA (green). **b.** Average DPOAEs thresholds at P30 for the same mice. Data are presented as mean ± SEM.

### Viral delivery of GJB2 with GREs rescues cochlear morphology in knockout mice

We then injected an AAV9-PHP.B vector encoding GJB2 under GRE control of expression (AAV-GRE-mmGJB2.HA; **Fig. 1a**) into cochleas of *Gjb2^Pfl/Pfl^, Cre+* conditional knockout mice at P1. As with expression of mmGJB2.HA in wild-type cochleas (**Fig. 1g**), the HA-tagged GJB2 was detected in many cells, including Deiters’ cells, Hensen’s cells, Claudius cells and border cells; fibrocytes type I, II and V of the lateral wall; and fibrocytes of the spiral limbus (**Fig. 2d**, **Table 1**)—congruent with the expression of eGFP (**Fig. 1f**) under the same regulatory elements. Importantly, no GJB2.HA signal was detected in IHCs and OHCs (parvalbumin label, yellow). We observed that restricted expression of GJB2 using AAV-GRE-mmGJB2.HA prevented the degeneration of the organ of Corti typically seen in *Gjb2^Pfl/Pfl^, Cre+* mice at P30. However, these mice still failed to develop the tunnel of the organ of Corti. Cells of the GJB2 network— comprising the epithelium of the inner and outer sulcus and the root cells of the spiral prominence, fibrocytes of the spiral limbus and strial cells—were preserved (**Fig. 2d**). Hair cells, missing at P30 in untreated *Gjb2^Pfl/Pfl^, Cre+* mice, were present in the vector-treated cochlea (**Fig. 2d**). Despite the preservation of cochlear structure, restriction of GJB2 expression using GREs only slightly mitigated the hearing loss in *Gjb2^Pfl/Pfl^, Cre+* mice (**Fig. 3**): ABR recordings at P30 demonstrated a 10–15 dB SPL improvement.

### Expression of GJB2 almost fully restores hearing in a milder mouse model of DFNB1

Although incorporation of GREs produced rescue of hearing greater than that previously reported, it did not restore ABR thresholds to wild-type levels. We speculated that GJB2 has a critical developmental role as early as P0^18^, and that vector delivered at P1 might not produce GJB2 protein until P2 or P3. We therefore sought a milder model of DFNB1, in which GJB2 expression is not entirely abolished, and turned to a mouse line in which the coding sequence of the upstream *Gjb6* gene was replaced with a Neo selection cassette and β-galactosidase reporter^37^. These substitutions in *Gjb6* have the effect of reducing GJB2 protein levels^38–40^ presumably due to disruption of nearby *Gjb2* regulatory sequences, and the mice are referred to here as *Gjb2^Gjb6-^*.

Immunofluorescent labeling with antibodies for GJB6 in the normal control *Gjb2^Gjb6+^* cochlea revealed that, similar to *Gjb2, Gjb6* is widely expressed in the epithelial and connective tissue cells of the cochlea (**Fig. 4a**, **Table 1**). The GJB6 signal was detected in Deiters’, Hensen’s, and Claudius cells, as well as in inner and outer pillar cells, in border cells, and in the epithelium of both the inner and outer sulcus. As expected, no GJB6 signal was observed in IHCs or OHCs. Examination of the lateral wall showed that GJB6 expression was high in type I and V fibrocytes, moderate in type II fibrocytes, and absent in type III and IV fibrocytes. In the stria vascularis, GJB6 puncta were found in the basal cells. Additionally, the fibrocytes of the spiral limbus contained GJB6 puncta.

**Figure 4.**
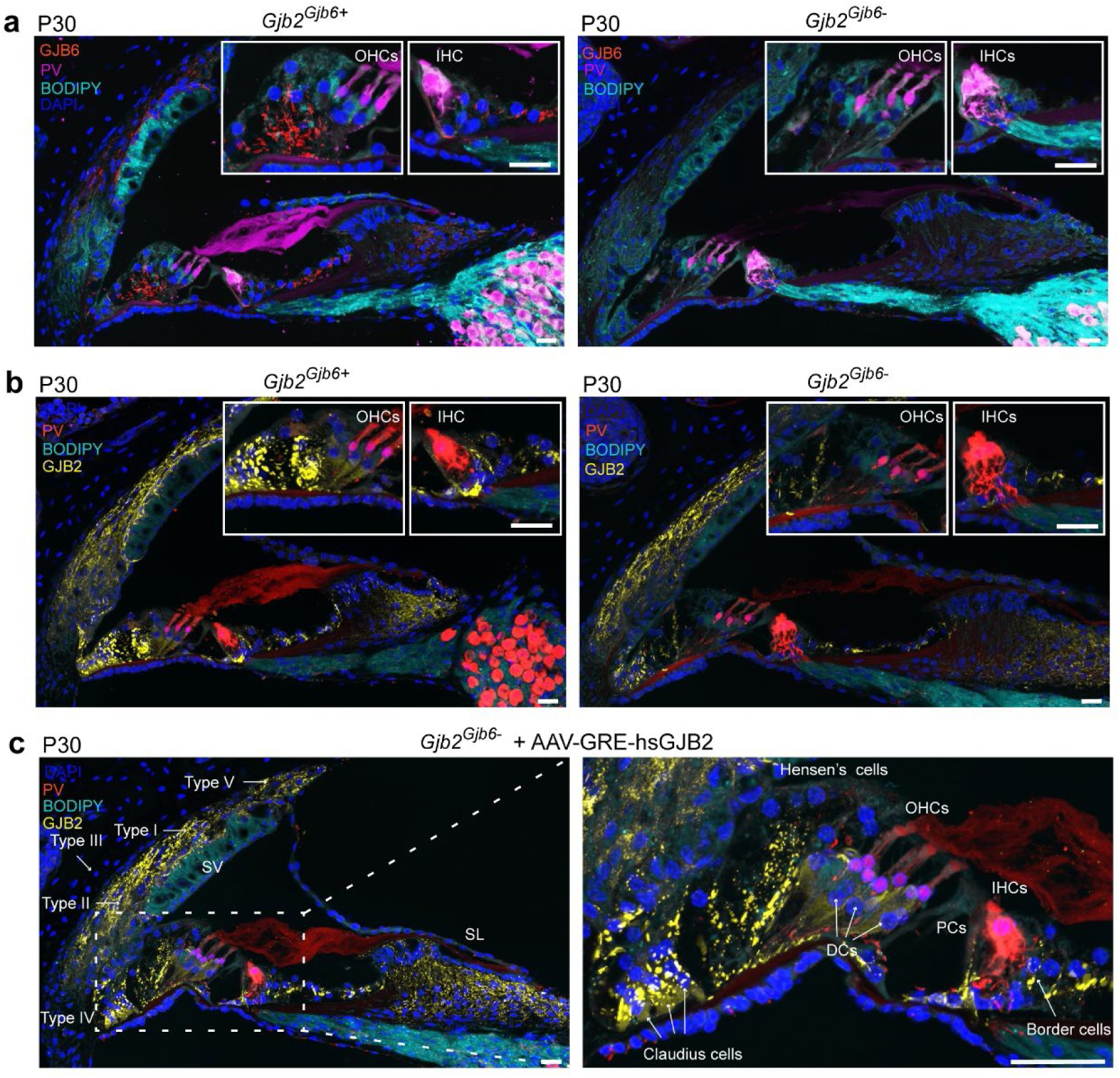
Rescue of pathology in *Gjb2^Gjb6-^* mice with AAV-GRE-hsGJB2. **a.** Representative confocal microscopy images for GJB6 in *Gjb2^Gjb6+^* control cochleas (*left*) and *Gjb2^Gjb6-^* mutants (*right*). Anti-GJB6 signal was detected in fibrocytes, Deiters’, Hensen’s, and Claudius cells, pillar cells and in both the inner and outer sulcus. No GJB6 immunoreactivity was observed in inner or outer hair cells. *Gjb2^Gjb6-^* cochleas showed normal gross morphology of the organ of Corti, hair cells, but no GJB6 was detected immunoreactivity (*right*). **b.** Representative confocal microscopy images of side-by-side comparison of GJB2 expression in *Gjb2^Gjb6-^* and *Gjb2^Gjb6+^* at P30. **c.** Representative confocal microscopy images of AAV- GRE-hsGJB2-treated *Gjb2^Gjb6-^* mice show complete restoration of GJB2 expression to levels seen in *Gjb2^Gjb6+^* controls. Scale bars: 20 μm.

Immunofluorescence analysis of *Gjb2^Gjb6-^* cochleas consistently demonstrated complete loss of GJB6 immunoreactivity (**Fig. 4a**). However, the mutant mice exhibited normal overall gross morphology of the organ of Corti, with normal presence of hair cells.

We observed a significant decrease in *Gjb2* expression in *Gjb2^Gjb6-^* mice compared to the *Gjb2* expression in *Gjb2^Gjb6+^* mice, particularly in Deiters’, Hensen’s, Claudius cells, and pillar cells, as well as in the epithelium of both the inner and outer sulcus (**Fig. 4b**, **Table 1**). However, *Gjb2* expression levels in *Gjb2^Gjb6-^* mice were similar to normal controls in the fibrocytes of the lateral wall and spiral limbus.

Since there were no observed differences in HA localization following the injection of AAV-GRE-mmGJB2.HA or AAV-GRE-hsGJB2.HA in wild-type control mice (**Fig. 1g, h**), indicating that mouse and human GJB2 exhibit similar localization and expression patterns in the mouse cochlea, we proceeded to inject *Gjb2^Gjb6-^* mice at P1 with AAV-GRE-hsGJB2.HA via the RWM. Thirty days post-injection, we analyzed cochlear morphology and hsGJB2.HA expression (**Fig. 5**). Despite strong HA labeling in cells that normally express GJB2, we observed several morphological abnormalities: a reduction in scala media volume, thickening of the tectorial membrane, enlargement of inner sulcus cells attached to the tectorial membrane, and a decrease or absence of the internal spiral sulcus. Additionally, there was a reduction or absence of the tunnel of Corti compared to untreated *Gjb2^Gjb6-^* and *Gjb2^Gjb6+^* mice. While IHCs appeared largely normal, OHCs were lost. This finding suggests that, despite strong expression and proper trafficking to the cell membrane, the GJB2 carrying the HA tag may not function properly.

**Figure 5.**
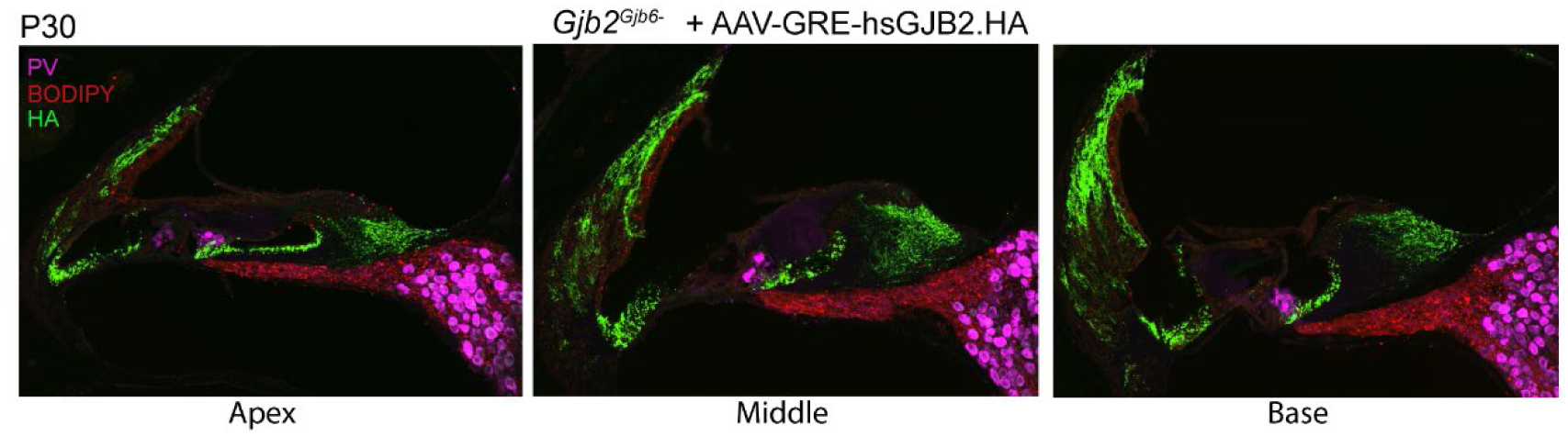
Morphological abnormalities in *Gjb2^Gjb6-^* mice injected with AAV-GRE-hsGJB2.HA. HA- tagged GJB2 expression 30 days post-injection. Despite strong labeling for hsGJB2.HA, multiple abnormalities were detected: reduced scala media volume, thickened tectorial membrane, enlarged inner sulcus cells, absent or reduced tunnel of Corti, and loss of OHCs.

Next, we injected *Gjb2^Gjb6-^* mice at P1 with AAV-GRE-hsGJB2, a vector encoding GJB2 without the HA tag. Thirty days after injection, the treated *Gjb2^Gjb6-^* mice receiving AAV-GRE-hsGJB2, showed a GJB2 network completely restored to levels observed in *Gjb2^Gjb6-^* control animals, and showed no histological abnormalities (**Fig. 4c**, **Table 1**). Apparently, HA-tagged GJB2 disrupts normal GJB2 function in this mouse model.

### AAV-GRE-hsGJB2 rescue hearing in Gjb2^Gjb-^ mice

We also assessed hearing function in untreated *Gjb2^Gjb6-^* and *Gjb2^Gjb6+^* mice, as well as in treated animals with AAV-GRE-hsGJB2.HA or AAV-GRE-hsGJB2. While *Gjb2^Gjb6+^* mice exhibited normal hearing, *Gjb2^Gjb6-^* mice were profoundly deaf by P30 (**Fig. 6a**). Next, we measured ABRs in response to broadband clicks and tone bursts, as well as DPOAEs in *Gjb2^Gjb6-^* mice injected with AAV-GRE-hsGJB2.HA. As expected, these mice showed no improvement in hearing, as tested by ABR or DPOAE (**Fig. 6a**). In contrast, when control mice were injected with AAV-GRE- hsGJB2.HA, their ABR results were similar to those of untreated control mice, and DPOAE levels were slightly elevated (**Fig. 6b**). These findings suggest that GJB2.HA is not toxic but is functionally inactive.

**Figure 6.**
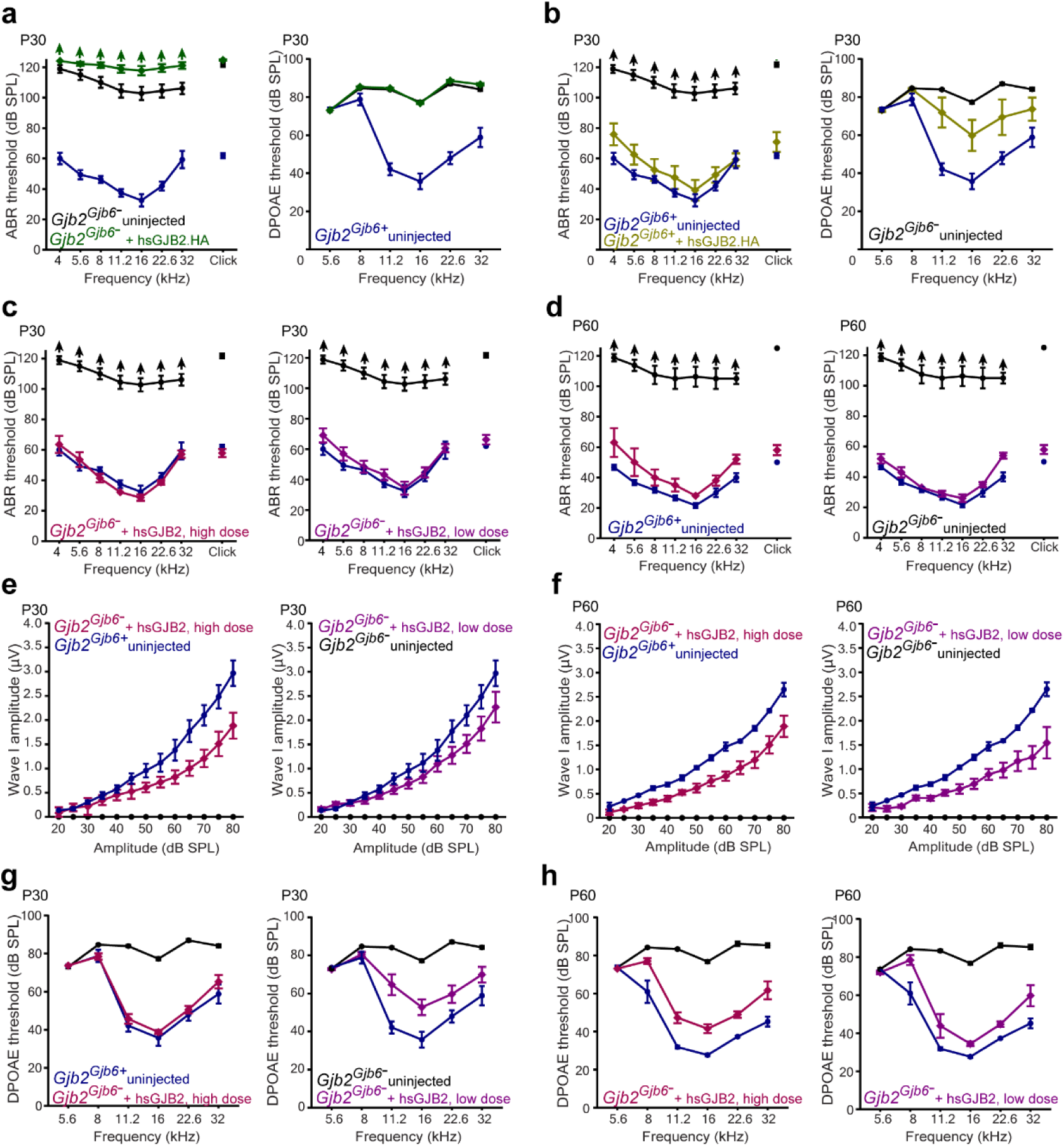
AAV-GRE-hsGJB2 rescues hearing in the *Gjb2^Gjb6-^* mouse model of DFNB1. **a.** Untreated *Gjb2^Gjb6+^* mice displayed normal hearing, while untreated *Gjb2^Gjb6-^* mice were profoundly deaf by P30. Hearing function was also evaluated in *Gjb2^Gjb6-^* mice treated with AAV-GRE-hsGJB2.HA. ABRs were measured in response to broadband clicks and tone bursts (left), alongside DPOAEs (right). *Gjb2^Gjb6-^* mice injected with AAV-GRE-hsGJB2.HA showed no improvement in hearing. **b.** Control mice injected with AAV- GRE-hsGJB2.HA displayed ABR responses comparable to untreated control mice. However, their DPOAE levels were slightly elevated. **c.** At P30, *Gjb2^Gjb6-^* mice treated with either a high dose (1.4×10¹¹ VGC) or low dose (7.0×10¹⁰ VGC) of AAV-GRE-hsGJB2 demonstrated full recovery of click-evoked and tone burst ABR responses, matching the levels observed in untreated control mice. **d.** Hearing recovery persisted through P60, with treated mice maintaining ABR responses equivalent to those of normal hearing controls. At P30 in **e** and P60 in **d**, the amplitude of ABR wave 1 (from P1 to N1) at 16 kHz in low-dose and high- dose treated mice. **g, h.** DPOAE measurements in *Gjb2^Gjb6-^* mice treated with AAV-GRE-hsGJB2. At P30 (**g**), DPOAE measurements demonstrated robust rescue in mice treated at P1 with either the high or low dose of AAV-GRE-hsGJB2. This rescue persisted at P60 (**h**), with DPOAE levels comparable to normal hearing mice.

Next, we evaluated hearing in mice injected with AAV-GRE-hsGJB2 at high (1.4×10^11^ VGC) and low (7.0×10^10^ VGC) doses. At P30, *Gjb2^Gjb6-^* mice treated with either dose demonstrated complete rescue of click-evoked and tone burst ABR responses to levels comparable to untreated control mice (**Fig. 6c**). This rescue persisted at P60 (**Fig. 6d**), with hearing comparable to that of normal hearing mice. Additionally, we analyzed the amplitude of ABR wave 1 at 16 kHz. At P30, the wave 1 amplitude in low-dose injected mice showed no significant difference from normal control mice. Similarly, in high-dose injected mice, it remained comparable to controls up to 55 dB SPL (**Fig. 6e**). High amplitudes were preserved up to P60 in mice expressing hsGJB2 (**Fig. 6f**). Measurements of DPOAEs showed robust rescue in mice treated at P1 with either dose at P30 (**Fig. 6g**) and P60 (**Fig. 6h**).

### Expression of GJB2 in nonhuman primate cochlea is similar to that in mouse

To assess viral delivery of GJB2 to the NHP cochlea, we first characterized the normal *GJB2* expression. We labeled frozen sections of vehicle-injected NHP cochlea with anti-GJB2 antibodies, and co-labeled supporting and hair cells with antibodies to SOX2 and MYO7A, respectively (**Fig. 7c**). As expected, multiplex labeling of GJB2 showed that protein is generally present in the epithelial and connective tissue cells. GJB2 signal was detected in Deiters’, Hensen’s, and Claudius cells, inner and outer pillar cells, and border cells (**Table 1**). Importantly, there was no GJB2 signal in IHCs or OHCs. A well-developed GJB2 network was also seen in the epithelium of the inner and outer sulcus and the root cells of spiral prominence. Analysis of the lateral wall revealed that *GJB2* was highly expressed in type I, II and V fibrocytes but was not present in type III and IV fibrocytes. In the stria vascularis, many GJB2 puncta were noted in the basal and intermediate cells but were not present in the marginal cells. The spiral limbus fibrocytes contained many GJB2 puncta, and many were present in the cells lining the scala vestibuli floor. We almost never saw GJB2 connecting interdental cells of the spiral limbus. The epithelium of Reissner’s membrane appeared devoid of GJB2. There was no *GJB2* expression in the modiolar region of the cochlea. However, we detected a very weak and diffuse signal in spiral SGNs, which we attribute to the autofluorescence of dense SGN cytoplasm. Comparison of all cochlear turns showed no gradient in *GJB2* expression from apex to base in the cells that express it.

**Figure 7.**
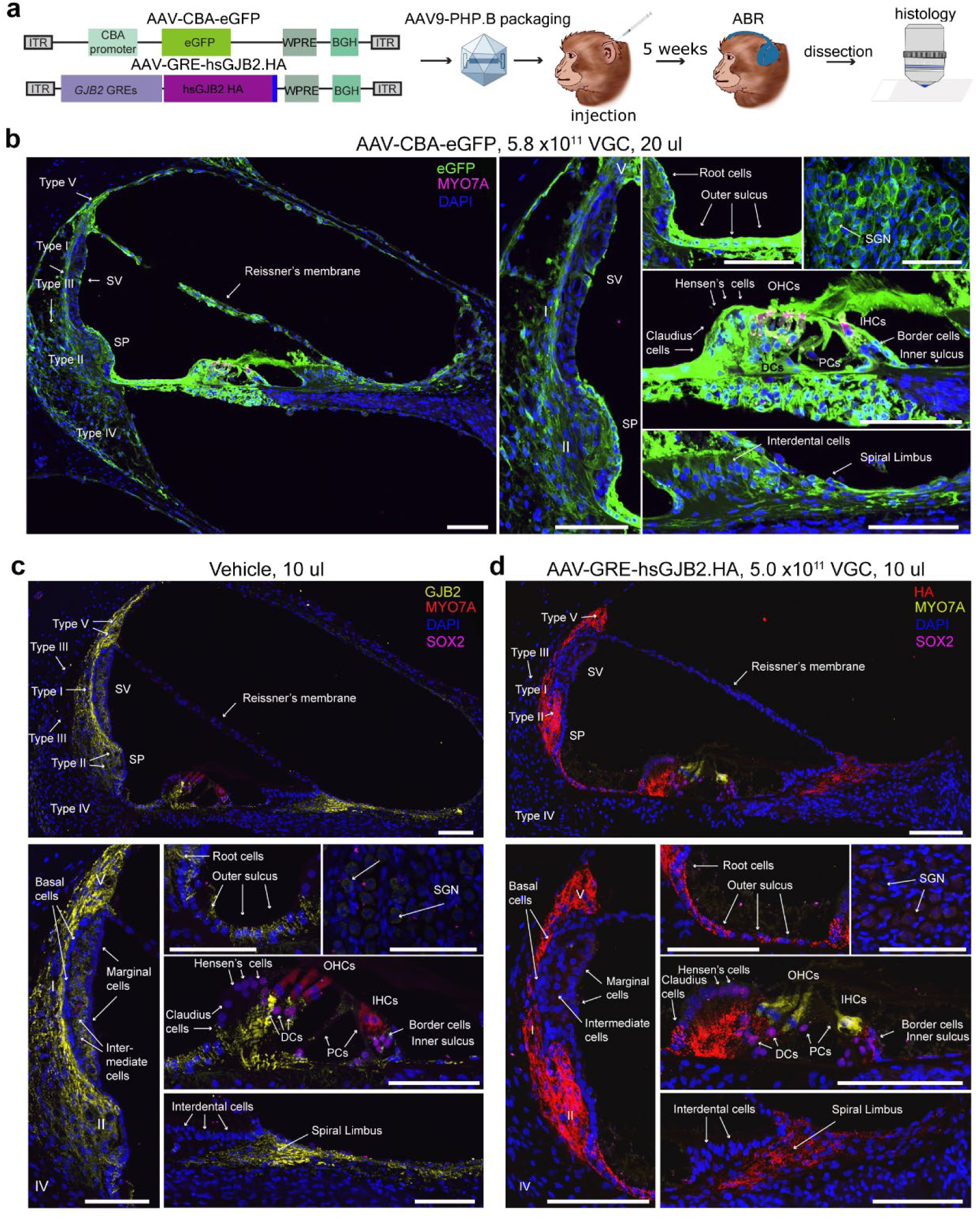
Expression of GJB2 by AAV-GRE-hsGJB2.HA is similar to that of native GJB2 in the NHP cochlea. **a.** Overview of the experimental design. **b.** Robust eGFP expression was observed in an NHP cochlea injected with AAV-CBA-eGFP. eGFP signal (green) was detected in hair cells, Deiters’ cells, Hensen’s cells, Claudius cells, pillar cells, border cells, the epithelium of the inner and outer sulcus and the root cells of the spiral prominence, in fibrocytes type I-V in the lateral wall and in the cells of the stria vascularis, in spiral limbus fibrocytes and in interdental cells. DAPI labeled nuclei (blue). **c.** Distribution of endogenous GJB2 (yellow) in an NHP cochlea. GJB2 signal was detected in the following supporting cells (magenta): Deiters’ cells (DCs), Hensen’s cells, Claudius cells, pillar cells (PCs), and border cells. The GJB2 network was seen in the epithelium of the inner and outer sulcus and the root cells of spiral prominence (SP), in fibrocytes type I, II and V in the lateral wall and in basal and intermediate cells in the stria vascularis (SV). The spiral limbus fibrocytes also contained many GJB2 puncta. Neither outer hair cells (OHCs) nor inner hair cells (IHCs) (red) expressed GJB2. DAPI labeled nuclei (blue). **d.** Exogenous (vector-delivered) GJB2.HA expression in an NHP cochlea. The HA tag was detected in SOX2-positive cells (magenta), such as in Deiters’ cells, Hensen’s cells, Claudius cells and border cells. Little or no HA labeling was seen in pillar cells. IHCs and OHCs (yellow) do not express GJB2.HA. The GJB2-HA network was seen in the epithelium of the inner and outer sulcus and the root cells of SP, in fibrocytes type I, II and V of the lateral wall and fibrocytes of the spiral limbus. The strial cells expressed exogenous GJB2.HA at a significantly low level. DAPI labeled nuclei (blue). Scale bars are 500 µm.

### GJB2 expressed in NHPs with a promiscuous CBA promoter is broadly localized

While functional rescue can be tested most easily in mice, larger animals such as nonhuman primates (NHPs) provide a more relevant model for transgene delivery to the human inner ear. We previously reported a successful surgical approach for delivering AAV vectors to the inner ears of cynomolgus monkeys^21–23^ and showed efficient transgene expression of a GFP reporter in a NHP cochlea following injection of the AAV9 capsid variant PHP.B via the RWM^21–23^. That vector used a ubiquitous CBA promoter, and we observed expression of the GFP reporter in many cell types, including off-target hair cells. Following our results in mouse, we expected more specific expression of GJB2 in NHP cochlea using GRE-driven viral vectors.

To directly compare cell-specific expression between CBA and GRE-driven constructs, we injected cochleas of three juvenile cynomolgus monkeys (*Macaca fascicularis)*. The study design is indicated in **Table 2** and **Fig. 7a**. The first animal received a vector encoding an eGFP marker under control of the CBA promoter by RMW injection in one ear (AAV-CBA-eGFP, 5.8×10^11^ VG, **Fig. 1b**). For the two other animals, three ears were injected with a vector encoding a hsGJB2.HA marker under the control of the GREs (AAV-GRE-hsGJB2.HA; 5.0×10^11^ VG, **Fig. 1b**); the fourth ear was injected with formulation buffer. Five weeks post-injection, animals were tested for ABR, then euthanized. Cochleas were extracted from the temporal bones of animals, further trimmed, decalcified and embedded in optimal cutting temperature (OCT) compound, cryosectioned and immunostained (**Fig. 7a)**.

**Table 2.**
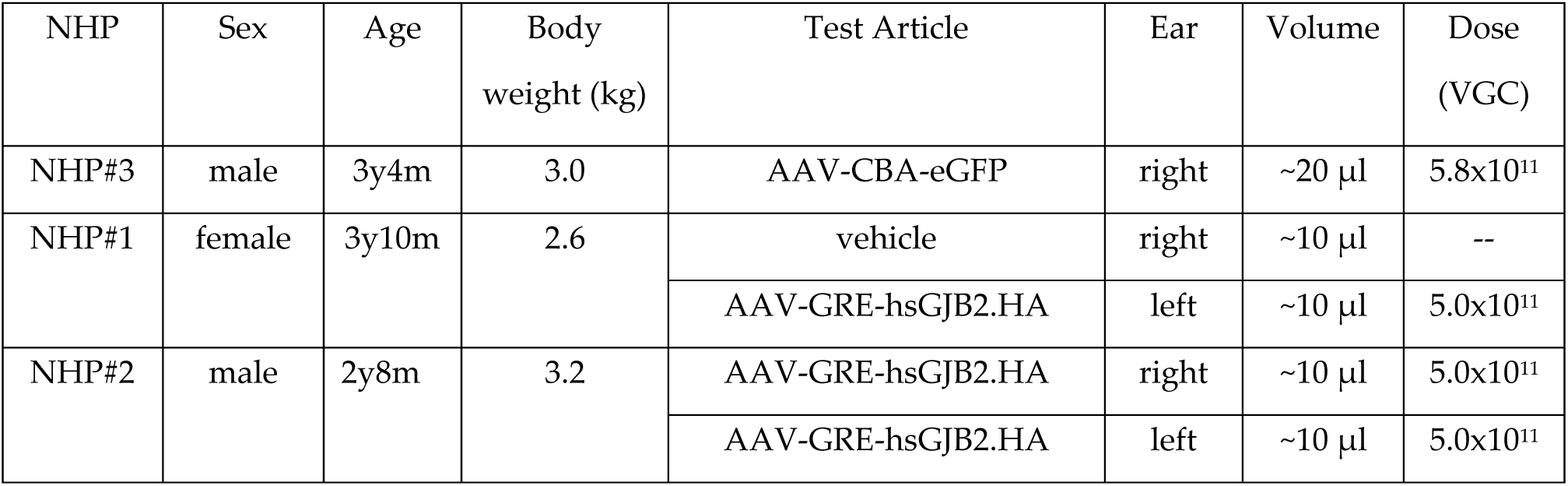
Study design.

Consistent with our results in mouse and our earlier NHP studies^21,22^, the AAV-CBA-eGFP vector showed wide-ranging transduction throughout the NHP organ of Corti, as revealed by anti-eGFP immunostaining of cochlea (**Fig. 7b**, **Table 1)**. Almost all cell types of the cochlea were transduced. In particular, we observed a very high level of eGFP expression in the organ of Corti, outer and inner sulcus cells, spiral limbus, lateral wall, and Reissner’s membrane. We analyzed higher-magnification images of the organ of Corti. In cochleas that received AAV-CBA-eGFP, complete transduction was observed in hair cells, Deiters’ cells, inner phalangeal cells, pillar cells, Hensen’s cells, and Claudius cells. Additionally, the epithelial cells of the inner sulcus and fibrocytes of the basilar membrane showed high transduction. Fibrocytes of the spiral ligament in the lateral wall were significantly transduced. Notably, high-magnification images of the modiolus region revealed high expression of eGFP in satellite glial cells, Schwann cells, and SGNs.

### GRE-driven viral GJB2 expression closely resembles endogenous expression in NHPs

In order to assess AAV-GRE-hsGJB2.HA expression in NHP cochleas, we first evaluated the specificity of the anti-HA antibody on frozen sections of vehicle-injected and AAV-GRE-hsGJB2.HA-injected NHP cochleas. We used antibodies to SOX2 and MYO7A to label supporting cells and hair cells, respectively. In the negative control vehicle-injected cochlea, as expected, we saw anti-SOX2 and MYO7A labeling but no HA signal (**Fig. 8a**). However, broad anti-HA labeling was detected in AAV-GRE-GJB2.HA-injected NHP cochleas (**Fig. 8b**). A weak and diffuse signal was detected in SGNs in both vehicle control and vector-injected cochleas, suggesting it represents SGN autofluorescence.

**Figure 8.**
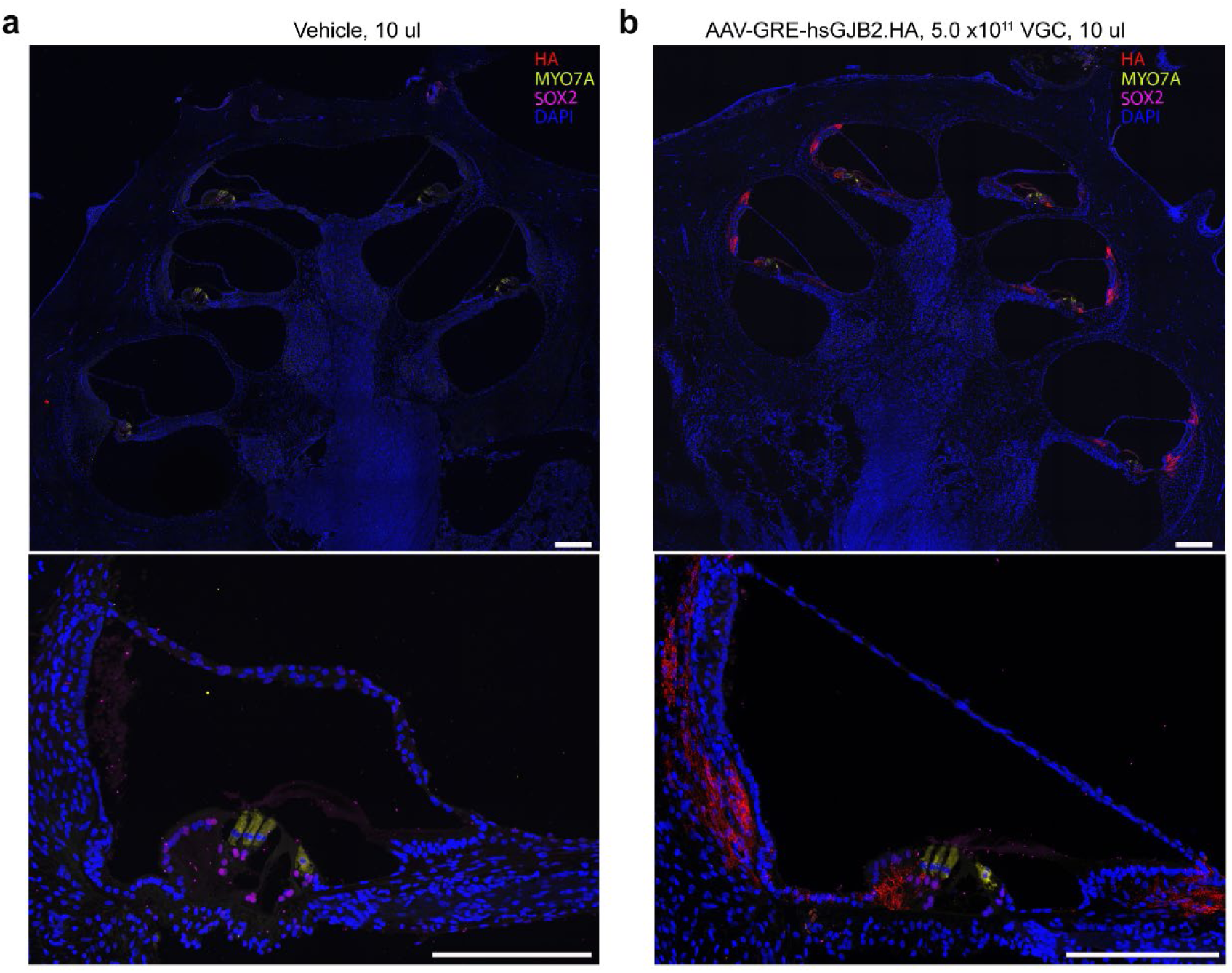
Uniform GJB2.HA expression across cochlear turns. **a.** Vehicle-injected cochlea. Anti-SOX2 labeling (magenta), anti-MYO7A labeling (yellow) and DAPI-labeled nuclei (blue) were detected. **b.** AAV- GRE-hsGJB2.HA-injected cochlea. Robust anti-HA labeling (red) was detected in the same locations as endogenous GJB2, analysis of cochlear turns showed no gradient in GJB2.HA expression between apex, middle and base. Scale bars = 1 mm.

Next, we investigated the virus-mediated GJB2.HA expression on cryosections collected from cochleas injected with AAV-GRE-hsGJB2.HA (**Fig. 7d**). Because injected cochleas have both endogenous and vector-delivered GJB2, we identified vector-delivered GJB2 with an antibody to the HA tag. With GJB2 expression driven by the *GJB2* GREs, IHCs and OHCs did not express hsGJB2.HA, as evidenced by lack of anti-HA labeling (**Fig. 7d**). Similar to the normal expression of GJB2 (**Fig. 7c)**, the anti-HA signal was detected in SOX2-positive cells, such as Deiters’ cells, Hensen’s cells, Claudius cells and border cells, with intensity similar to that of native GJB2 (**Table 1**). However, little or no HA-tagged puncta were seen in inner or outer pillar cells. A well-developed exogenous GJB2.HA network was also seen in the inner and outer sulcus epithelia and the root cells of spiral prominence. The spiral limbus fibrocytes and the cells lining the scala vestibuli floor expressed GJB2.HA in a pattern similar to that of native GJB2. We detected weak or no HA signal in the spiral limbus interdental cells. Analysis of the lateral wall showed that, like native GJB2, exogenous GJB2.HA was highly expressed in type I, II and V fibrocytes and was not present in type III and IV fibrocytes. As expected, there was also no expression of GJB2.HA in the marginal cells or the epithelium of Reissner’s membrane. There was no GJB2 expression in the modiolar region of the cochlea. As with endogenous GJB2, analysis of cochlear turns showed no gradient in GJB2.HA expression between apex, middle and base (**Fig. 8b**). There were some differences: the stria vascularis and pillar cells expressed exogenous GJB2.HA at lower level than endogenous GJB2.

### AAV-GRE-hsGJB2.HA is safe in NHP cochleas

We performed histological analysis on H&E-stained frozen tissue sections from injected cochleas to evaluate pathology (**Fig. 9a**). Tissue morphology and architecture appeared normal in both AAV-treated and vehicle control groups, with no significant abnormalities observed. Minor changes in the basal turn, likely related to surgery and vector injection, were noted in the left cochlea of NHP#2. To further evaluate immune response activation in the cochlea we performed staining against ionized calcium-binding protein (IBA1) in frozen tissue sections from injected and control cochleas (**Fig. 9b**). IBA1 is primarily expressed in microglia cells, which in the cochlea are macrophages. Macrophages are abundant in the cochlea; however, insults such as AAV capsids or surgery can cause an inflammatory response, which includes activation of tissue-resident macrophages as characterized by changes in macrophage morphology, number, and distribution. We observed that normal tissue morphology and architecture appeared to be preserved in all cochleas, with no notable abnormalities. (**Fig. 9b**). Sensory hair cells and supporting cells appeared normal. We observed no difference in macrophage infiltration in treated cochleas compared to vehicle-injected cochlea, confirming that NHPs injected with AAV-GRE-hsGJB2.HA showed minimal immune response to the vector.

**Figure 9.**
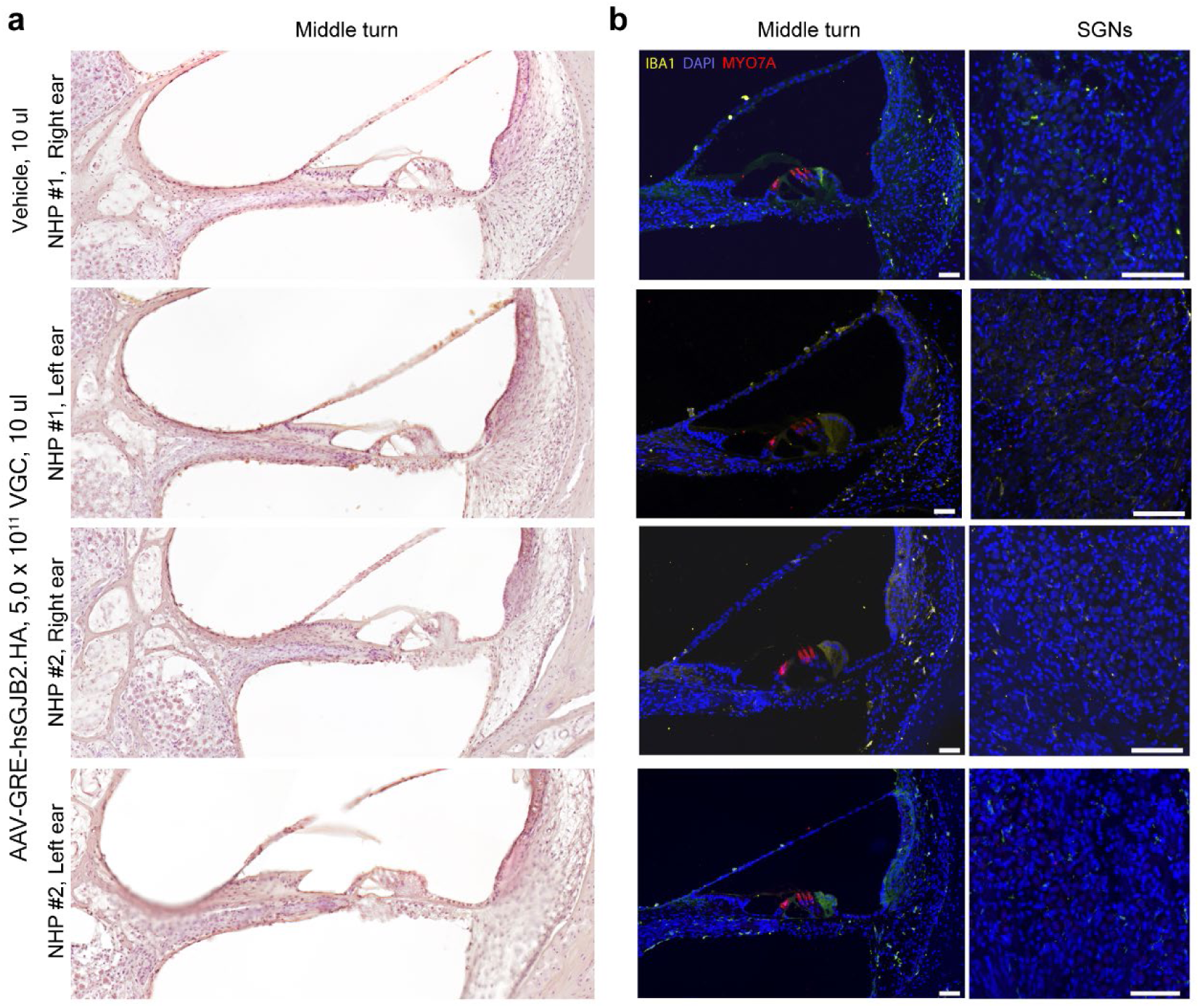
Histological evaluation of safety. **a.** Hematoxylin and eosin-stained sections of NHP cochlea injected with AAV-GRE-hsGJB2.HA 5.0×10^11^ VGC. Tissue morphology and architecture appeared normal in both AAV-treated and vehicle control groups, with no significant abnormalities observed. **b.** No immune infiltration in the inner ear compared to vehicle-injected cochlea, confirmed by labeling of ionized calcium-binding protein (IBA1) (yellow). No abnormalities were seen in hair cells (red). DAPI labeled nuclei (blue). Scale bars = 500 µm.

To assess safety physiologically, we also tested the hearing before and after injection with either AAV-GRE-hsGJB2.HA or vehicle (**Fig. 10)**. We compared pre-injection thresholds to thresholds at 5 weeks after injection to assess changes that may be due to the vector. In the vehicle-injected cochlea (NHP#1, right ear), no significant difference was seen in the click and the pure tone ABR threshold after injection. In two ears injected with 5.0× 10^11^ VG (NHP#1 left ear and NHP#2 right ear), we saw no significant change in sensitivity in either click or tone ABR. However, in the one ear injected with 5.0×10^11^ VG (NHP#2 left ear), hearing threshold was slightly elevated at all frequencies by 15-25 dB nHL. This may be a result of early postoperative changes in the middle ear after mastoid surgery. Because two of three ears showed no elevation of threshold, we believe the injection of vector does not reproducibly cause a loss of sensitivity.

**Figure 10.**
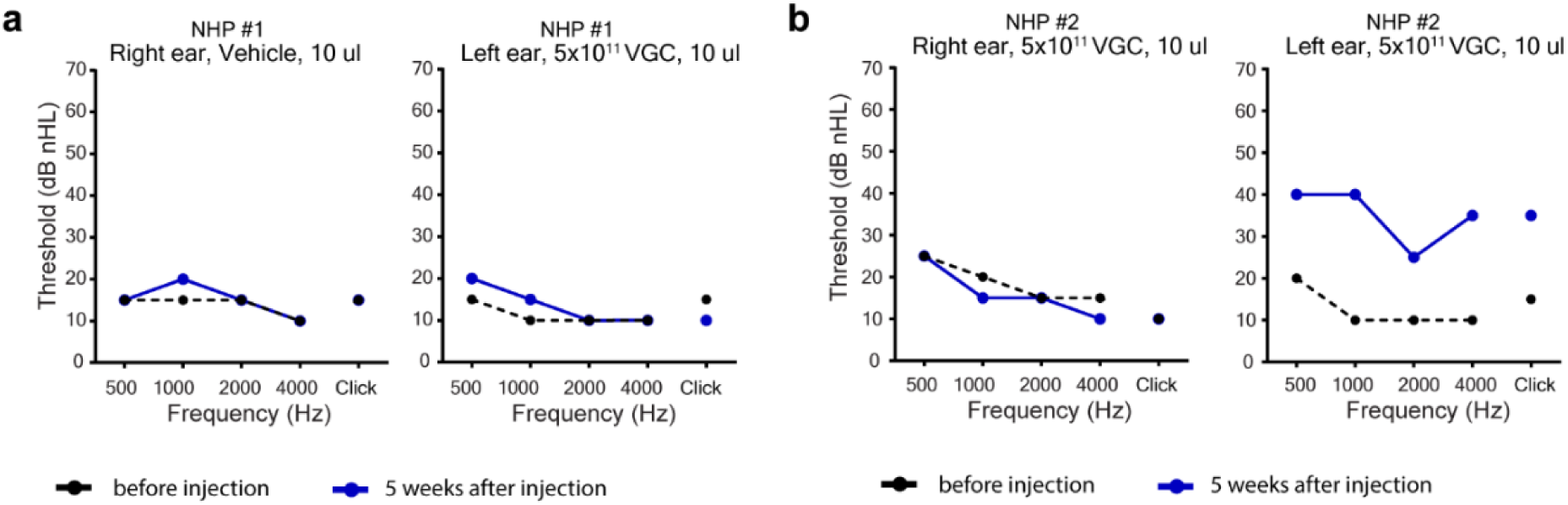
Physiological tests of safety. **a, b.** NHP #1 (**a)** and NHP #2 (**b**) were tested with click and tone ABR before AAV-GRE-hsGJB2.HA or vehicle control injection (black) and then five weeks later, before euthanasia (blue).

## Discussion

Studies on gene therapy for deafness typically use AAV stereotypes that transduce many cell types in the inner ear, including cell types that do not express the gene being delivered, and they often use strong ubiquitous promoters to drive gene expression. This approach has successfully restored hearing and balance in several other mouse models of human deafness, without documented toxicity^21,23,41–55^. In the case of GJB2, however, overexpression and mis-expression led to toxicity ranging from increased hearing thresholds to lethality^17,19,20^, suggesting a need for more specific vector design.

One way to restrict expression is by choosing an AAV serotype that targets a cell type of interest. In the case of *GJB2*, there are no reported AAV serotypes that mimic its expression pattern. Moreover, *GJB2* is expressed in both epithelial cells and fibrocytes of the cochlea, and it is unlikely that one AAV capsid would transduce those two cell types and no others. Instead, we chose to target expression by choosing a capsid, AAV9-PHP.B, that transduces all the cells of interest, and to limit gene expression by harnessing cis-acting gene regulatory sequences. To identify candidate GREs governing *Gjb2* expression in the mouse cochlea, we performed ATAC-seq on neonatal cochleas at P2, P5, and P8, identifying multiple regions of open chromatin near the mouse *Gjb2* gene. We prioritized GREs by avoiding neuronal-associated regions and focusing on elements conserved across mammalian genomes.

We first tested the selected GREs in a vector driving expression of eGFP. In injected cochleas at P6, eGFP expression overlapped normal *Gjb2* expression and was eliminated from the cochlear hair cells. Next, we replaced the eGFP coding sequence with that for mmGJB2.HA (AAV-GRE-mmGJB2.HA) or hsGJB2-HA (AAV-GRE-hsGJB2.HA). Neonatal wild-type cochleas injected with mmGJB2.HA or hsGJB2.HA vectors exhibited similar HA-tag expression at P30, consistent with normal GJB2 localization and proper membrane trafficking, confirming that the GREs effectively restricted expression. Injection of AAV-GRE-mmGJB2.HA into the *Gjb2^Pfl/Pfl^, Cre+* mouse model for DFNB1 preserved cochlear morphology and improved hearing thresholds by 10–15 dB SPL at P30.

ABR thresholds did not reach wild-type levels, however, perhaps because GJB2 protein is needed for development before the vector injection at P1. To address this possibility, we used a milder DFNB1 model, *Gjb2^Gjb6-^*, in which GJB2 protein levels are reduced but not entirely abolished due to putative disruptions of *Gjb2* regulatory sequences^38–40^ and ABR thresholds are severely but not profoundly elevated. Immunofluorescence analysis of untreated *Gjb2^Gjb6-^* cochleas showed no GJB6 immunoreactivity but retained normal organ of Corti morphology with intact hair cells.

Injection of AAV-GRE-hsGJB2.HA into *Gjb2^Gjb6-^* mice resulted in strong HA-tag expression in cells that normally express GJB2, but structural abnormalities were observed, including scala media volume reduction, tectorial membrane thickening, and OHC loss, with the phenotype being more severe than in uninjected mutant mice. This finding suggests that when GJB2 is partially functional (GJB2.HA) and GJB6 is absent, it leads to detrimental changes in the inner ear. No such changes were observed in *Gjb2^Gjb6+^* control mice after AAV-GRE-hsGJB2.HA injection, which displayed normal ABR and morphology.

We injected *Gjb2^Gjb6-^* mutant mice at P1 with AAV-GRE-hsGJB2, in which GJB2 lacked the HA tag. Thirty days later, immunofluorescence analysis showed complete restoration of the GJB2 network with no histological abnormalities. Hearing assessments revealed that both high and low doses of the vector led to complete rescue of ABR responses at P30, with the effects persisting to at least P60. GREs apparently restrict expression to only the cells that express GJB2, GJB2 traffics to the cell membrane to form gap junctions, and only untagged GJB2 results in fully functional gap junctions.

While testing functional rescue is more straightforward in mice, larger animals like NHPs serve as a more appropriate model for transgene delivery to the human inner ear. Here, we also report for the first time expression of GJB2.HA under cis-acting *GJB2* regulatory sequences in an NHP inner ear. With AAV-mediated GJB2.HA expression in NHP cochleas, we detected vector-delivered GJB2 in SOX2-positive cells, including Deiters’ cells, Hensen’s cells, and fibrocytes, but not in IHCs or OHCs. The expression pattern of exogenous GJB2.HA was similar to that of native GJB2 in most cochlear regions (**Table 1**).

For both the mouse and the NHP, we chose human cis-regulatory sequences. Their efficacy preserving morphology and hearing sensitivity, and lack of toxicity in both the control mice and NHPs, suggest we have captured at least some conserved sequences required to mediate appropriate *GJB2* expression in the human inner ear. Further work is needed to determine the minimum sequence required to provide a therapeutic effect without toxicity.

Taken together, our data show the importance of targeting a therapeutic vector to the correct cells. In this study we regulated gene expression by employing native cis-acting regulatory sequences, rather than cell-type-specific AAV capsids. For the first time, we demonstrate substantial hearing rescue in a pre-clinical model of human DFNB1 hearing loss and show that in wild-type NHPs, GREs effectively restrict GJB2 expression to the correct cells, confirming both their precision and safety. This should be an important step towards developing a gene therapy for human GJB2-related hearing loss.

## Methods

### Study approval

All animal studies were conducted in compliance with ethical regulations according to protocol IS00003633 approved by the Institutional Animal Care and Use Committee (IACUC) at Harvard Medical School, Boston, and were performed according to the NIH guidelines.

### Mouse models

*Gjb2^Pfl^* conditional knockout mice (*Gjb2^tm1Ugds^*)^34^ have loxP sites flanking the coding sequence in exon 2. *Gjb2^Pfl^* mice were crossed to *Sox10-Cre* mice (*Tg(Sox10-cre)1Wdr/J*, Jackson Laboratory #025807)^56^, which have a BAC transgene encoding Cre recombinase under the *Sox10* promoter. *Gjb6^-^* constitutive knockout mice (*Gjb6^tm1Kwi^*; European Mouse Mutant Archive #EM:00323)^37^ have the *Gjb6* coding region replaced by a Neo cassette and β-galactosidase reporter with a nuclear localization sequence. These substitutions have the effect of reducing GJB2 protein levels^38–40^, so are referred to here as *Gjb2^Gjb6-^*.

### AAV construction

Plasmids for AAV production were constructed as previously described^21,22^. The CBA promoter consisted of the cytomegalovirus (CMV) early enhancer element; the promoter, first exon, and first intron of the chicken beta-actin gene; and the splice acceptor of the rabbit β-globin gene. Candidate GREs were identified through ATAC-seq. Vectors included a woodchuck hepatitis virus post-transcriptional regulatory element (WPRE) and a bovine growth hormone poly A signal (BGH).

For all experiments, the capsid was AAV9-PHP.B^28^. AAV packaging was performed by the Boston Children’s Hospital Viral Core (Boston, MA, USA) or by PackGene Biotech (Houston, TX, USA). All plasmids were sequence-verified prior to packaging. Viral titers were determined by qPCR specific for the inverted terminal repeat of the virus.

### ATAC-seq library preparation and sequencing

Neonatal (P2, n=3; P5, n=3; P8, n=4) CD1 mice were sacrificed and cochleas were dissected bilaterally as described above prior to fixation. All tissue from each animal was combined and dounced 10 times in 5 ml of HB buffer (0.25 M sucrose, 25 mM KCl, 5 mM MgCl_2_, 20 mM Tricine-KOH pH 7.8, 1 mM DTT, 0.15 mM spermine, 0.5 mM spermidine, DTT 1 mM [Sigma], half of a Roche proteinase inhibitor tablet) in a 7 ml dounce-homogenizer via tight pestle, then supplemented with IGEPAL CA-630 to final concentration of 0.3%, and finally dounced an additional five times with the tight pestle. Nuclei were filtered through a 40 μm strainer and gently combined 1:1 with 50% iodixanol. Nuclei were then underlaid with a gradient of 1 ml 40% iodixanol and 1 ml 30% iodixanol and centrifuged at 10,000*g* for 19 minutes using the “no brake” setting. Purified nuclei (0.8 ml) were then collected from the 30/40% interface. A 20 μL aliquot was taken for visual inspection and cell counting, with typical yield of ∼100,000-200,000 cochlear nuclei per animal.

ATAC-seq (assay for transposase accessible chromatin with sequencing) was performed using the OMNI-ATAC protocol^57^ immediately following the nuclear isolation steps described above. Nuclei (50,000) were pelleted at 500 *g* for 10 minutes at 4°C in a fixed angle centrifuge and resuspended in 50 μL transposition buffer (prepared from Illumina kit components TD buffer and 100 nM transposase (Illumina), as well as 0.01% digitonin, 0.1% Tween-20) via careful pipetting. Transposition occurred at 37°C in a thermomixer at 1000 rpm for 30 minutes before resuspension in 250 μL of DNA binding buffer (Zymo DNA Clean and Concentrator Kit) and reaction cleanup per manufacture instructions, with final elution volume of 21 μL in elution buffer. Samples were amplified following NEBNext High-Fidelity 2X PCR Master Mix manufacturer protocols (NEB) in a total reaction volume of 50 μL (1.25 μM each of i5 and i7 sequencing primers) at 72°C for 5 minutes, 98°C for 30 seconds, (98°C for 10 seconds, 63°C for 30 seconds, and 72°C for 1 minute) x10 cycles. PCR samples were re-purified following the Zymo manufacturer protocol and eluted into 21 μL of elution buffer. Size selection was performed via agarose gel (150-1000 bp) followed by Qiagen MinElute gel extraction according to the manufacturer’s instructions, with two serial 10 μL elution steps. Seventy-five bp paired-end sequencing was performed on an Illumina NextSeq500.

### ATAC-seq peak calling and quantification

Putative cochlear GREs (regions of ATAC-seq enrichment) were identified as described previously^32^. Briefly, peaks were determined using MACS2 (v 2.1.0) parameters -p 1e-5– nolambda –keep-dup all –slocal 10000, as previously described^29^. Peaks from individual sample replicates were intersected to find only reproducible regions of enrichment. Blacklist regions were removed as previously described^58^. To identify sites with enriched ATAC-seq signal (peaks), we applied the IDR pipeline using the MACS2 peak calling algorithm^59^ with the following parameters: –nomodel –extsize 200 –keep-dup all. An IDR threshold of 0.01 was used for self-consistency, true replicate, and pooled-consistency analyses. The ‘optThresh’ cutoff was then used to obtain a final set of high-confidence, reproducible ATAC-seq peaks for each sample.

To produce a final list of reference coordinates of all genomic regions that were accessible in at least one murine cochlear ATAC-seq sample, the MACS2-called peaks for each experimental replicate were unioned using the bedops --everything command. Bedtools merge was then used to combine any peaks that overlapped in this unioned bed file; in this way, any region that was significantly called a peak in at least one ATAC-seq dataset was incorporated in the final aggregated peak list. The featureCounts package was then used to obtain ATAC-seq read counts for each of these accessible putative CREs for downstream analyses^60^, including comparison to previously published murine neuronal ATAC-seq data^61^ to identify putative cochlear-restricted sites.

### AAV round window membrane injection in neonatal mice

RWM injections were performed under a stereomicroscope (Nikon SMZ1500). P0-P1 pups were anesthetized using cryoanesthesia and kept on an ice pack during the procedure^21,23,44^. A small incision was made beneath the external earbud, which was then extended to allow for separation of the soft tissues and exposure of the bulla. The round window niche was identified visually. Using a Nanoliter 2000 Injector (World Precision Instruments), 1 μl of the viral vector solution was injected through a micropipette needle. The incision was closed with a 7-0 Vicryl surgical suture, and standard postoperative care was provided following the injection.

### Mouse cochlear histology and imaging

In experiments where transduction efficiency of AAV-eGFP was detected via its intrinsic fluorescence, organ of Corti explants were dissected at P6 in L-15 medium and fixed with 4% formaldehyde in Hank’s balanced salt solution (HBSS) for 1 h, washed three times with HBSS, and then blocked and permeabilized with 10% donkey serum with 0.3% Triton X-100 for 1 h at room temperature. Rabbit anti-GJB2 antibody (1:100) (#71-0500, Fisher) was used to label GJB2. Antibody was diluted in 10% donkey serum and incubated overnight at room temperature followed by several washes in HBSS. Next, samples were incubated in blocking solution for 30 min and incubated overnight at room temperature with a donkey anti-rabbit secondary antibody conjugated to Alexa Fluor 593 in a 1:500 dilution in blocking solution. To label hair bundle actin, we used phalloidin conjugated to Alexa Fluor 405 (1:20; Life Technologies).

In experiments assessed with cryosections, mouse cochleas were dissected in L-15 medium, immediately fixed with 4% formaldehyde in HBSS for 1 h at room temperature, then washed with HBSS and transferred to fresh 10% EDTA for 1-3 days. Next, samples were cryoprotected and embedded in OCT compound and stored at −80°C prior to sectioning. Cryosections were generated using a Leica CM 3050 S cryostat at 20 μm step size.

For immunofluorescence labeling, the following primary antibodies and secondary antibodies were used: rabbit anti-HA C29F4 antibody (1:200) (#3724, Cell Signaling), rabbit anti-GJB2 antibody (1:50) (#71-0500, Fisher), anti-parvalbumin antibody (1:200) (#195004, Synaptic System), rabbit anti-GJB6 antibody (1:200) (#71-2200, Fisher), rabbit anti-MYO7A (1:200) (#25-6790; Proteus Biosciences), donkey anti-rabbit IgG secondary antibody conjugated to Alexa Fluor 594 (1:200), and donkey anti-rabbit IgG conjugated to Alexa Fluor 488 (1:200), donkey anti-guinea pig Alexa Fluor 647 (1:200). We used phalloidin conjugated to Alexa Fluor 405 (1:20; Life Technologies) to label hair-bundle actin (1:20), DAPI-labeled nuclei (1:500), BODIPY (4,4-difluoro-4-bora-3a,4a-diaza-*s*-indacene; Invitrogen, D3835) to label membranes (1:1500). Samples were blocked for 1 h at room temperature. Antibodies were diluted in 10% donkey serum and incubated overnight at room temperature, followed by several rinses in HBSS. Next, samples were incubated in a blocking solution for 30 min and incubated overnight at room temperature with a secondary antibody in the blocking solution. Tissues were mounted on a Colorfrost glass slide (Thermo Fisher Scientific) using Prolong Gold Antifade mounting medium (Thermo Fisher Scientific). Imaging was performed with a Nikon Ti2 inverted spinning disk confocal using a Plan Fluor 40×/1.3 oil objective, Plan Apo λ 60×/1.4 oil objective or Plan Apo λ 100×/1.45 oil objective.

### Scanning electron microscopy

SEM in neonatal and adult mice was performed as previously described^62^. Neonatal organ of Corti explants were dissected at P6 in L-15 medium and fixed with 2.5% glutaraldehyde in 0.1 M cacodylate buffer (pH 7.2), supplemented with 2 mM CaCl_2_ for 1–2 h at room temperature. The samples were rinsed three times in sodium cacodylate buffer, 0.1 M, pH 7.4 for 10 min, and then briefly rinsed once in distilled water, dehydrated in an ascending series of ethanol concentrations, and critical-point dried from liquid CO_2_ (Tousimis Autosamdri 815). Samples were mounted on aluminum stubs with carbon conductive tabs and were sputter-coated (EMS 300 T dual-head sputter coater) with platinum to 5 nm and observed in a field-emission scanning electron microscope (Hitachi S-4700).

### Mouse ABR and DPOAE testing

ABRs and DPOAEs were recorded as previously described^44,63^, utilizing an acoustic system developed by Massachusetts Eye and Ear, Boston, MA, USA. Briefly, adult mice were administered anesthesia and maintained on a 37°C heating pad throughout the recording session. Tone-pip stimuli (5 ms duration and 0.5 ms rise-fall time) were delivered at frequencies ranging from 4 kHz to 32 kHz. Sound levels were increased in 5-dB increments from 20 to 120 dB sound pressure level (SPL). ABR Peak Analysis software (version 1.1.1.9, Massachusetts Eye and Ear, Boston, MA, USA) was used to measure ABR thresholds and peak amplitudes. DPOAEs were recorded for primary tones with a frequency ratio of f2/f1=1.2, with L1=L2+10 dB. The f2 frequency ranged from 5.6 kHz to 32 kHz in half-octave increments. Primary tone levels were adjusted in 5-dB increments, from 10 to 120 dB SPL for f2.

### NHP study design

One female and two male juvenile cynomolgus monkeys (*Macaca fascicularis*; 2–4 years old, weighing 2.6–3.2 kg) were used in this study. Primate work was performed at Biomere Medical Research Models (Worcester, MA, USA) according to Biomere’s animal use guidelines and approved procedures. Each animal was considered acclimated to the environment at the time of the study. NHPs were healthy and without a history of ear inflammation, ear surgery, signs of balance disorders, or other risk factors for dysfunction of the inner or middle ear. Monkeys received transmastoid/trans-RWM injections of 10 μL of AAV-GRE-hsGJB2.HA vector with doses of 5.0 × 10^11^ VG (n = 3 ears) and 20 μL AAV-CBA-eGFP of 5.8 × 10^11^ VG (n = 1 ear). One animal was injected in one ear with 10 μL formulation buffer control (see Table 2). A hearing test was performed before, and 5 weeks after unilateral injection in NHP #1 and NHP#2. Five weeks post-injection, animals were euthanized. Cochleas were extracted from the temporal bones of animals for histological analysis.

### NHP surgical procedures

Pre- and post-operative anesthesia and analgesia were performed as described previously^22^. The animals were sedated with ketamine and pre-medicated with atropine following overnight food deprivation. Prior to surgery, the animals were administered dexamethasone and buprenorphine and given a single dose of sustained-release buprenorphine and cefazolin. During the surgery, they were maintained with oxygen and isoflurane. The surgical site was prepared for aseptic surgery, and the animals’ vital signs were monitored. Bupivacaine was used for local anesthesia.

All viral inner-ear administrations were performed via the facial recess approach, with round-window injection as described^22,23^. The viral vector was injected using a microinjection pump with a Hamilton syringe and a sharp-end stainless steel needle. After injection, the surgical site was flushed with warm saline and closed with tension and absorbable sutures. The skin was cleaned with saline and a topical antibiotic ointment was applied. The wounds were monitored for proper healing for 2 weeks.

### NHP cochlear processing, histology, and imaging

NHPs were euthanized at 35 days after vector injection and were perfused with heparinized saline followed by 4% formaldehyde. Collected temporal bones were postfixed for another 48 h in 4% formaldehyde. Then, cochleas were dissected from the temporal bones under a microscope as described^22,23^ and decalcified in EDTA. Next, samples were cryoprotected by incubating in gradient concentrations of sucrose and were embedded in OCT compound and stored at −80°C prior to sectioning. Cryosections were generated using a Leica CM 3050 S cryostat at 30-μm step size.

For immunofluorescence labeling, the following primary antibodies and secondary antibodies were used: mouse anti-MYO7A antibody (1:50) (#sc-74516, Santa Cruz), rabbit anti-HA C29F4 antibody (1:200) (#3724, Cell Signaling), rabbit anti-GJB2 antibody (1:50) (#71-0500, Fisher), goat anti-SOX2 antibody (1:50) (#AF2018, R&D Systems), rabbit anti-IBA1 antibody (1:200) (#019-19741, Wako Chemicals), donkey anti-rabbit IgG secondary antibody conjugated to Alexa Fluor 594 (1:200) (Invitrogen), donkey anti-goat IgG conjugated to Alexa Fluor 647 (1:200) (Invitrogen), donkey anti-mouse IgG conjugated to Alexa Fluor 488 (1:200) (Invitrogen), donkey anti-mouse IgG conjugated to Alexa Fluor 594 (1:200) (Invitrogen), and donkey anti-rabbit IgG conjugated to Alexa Fluor 488 (1:200) (Invitrogen). Samples were blocked for 1 h at room temperature. Antibodies were diluted in 10% donkey serum and incubated overnight at room temperature, followed by several rinses in HBSS. Next, samples were incubated in a blocking solution for 30 min and incubated overnight at room temperature with a secondary antibody in the blocking solution. We used DAPI to label cell nuclei. Tissues were mounted on Colorfrost glass slides (Thermo Fisher Scientific) using Prolong Gold Antifade mounting medium (Thermo

Fisher Scientific). Imaging was performed with a Nikon Ti2 inverted spinning disk confocal using a Plan Fluor 40×/1.3 oil objective and Plan Apo λ 60×/1.4 oil objective.

### ABR testing in NHPs

Testing was performed as described previously^23^. A hearing test was performed before, and 5 weeks after the unilateral injection. The monkeys were anesthetized with a mixture of ketamine and atropine and were masked with isoflurane. During the procedure, the animal was maintained on isoflurane anesthesia. ABRs were acquired using a clinical-grade auditory evoked potential system (GSI Audera v2.7; Grayson-Stadler, Eden Prairie, MN, USA). Scalp potentials were recorded using subdermal needle electrodes applied in a 3-electrode configuration. The click and tone-pip stimuli were presented at alternate polarity with a tubal insert phone (TIP 50 GSI generic; Grayson-Stadler, Eden Prairie, MN, USA). The acoustic click was presented at a repetition rate of 11.1 Hz, and waveforms were averaged across 2,004 stimulus repetitions. Clicks were a condensation stimulus of 100-μs duration. Sound levels were incremented in 5-dB steps, from ∼15 dB nHL up to 54.5 dB nHL. Tone frequencies of 500, 1000, 2000 and 4000 Hz were used. The tone pips were presented at a repetition rate of 27 Hz and waveforms were averaged across 2,016 stimulus repetitions. Sound levels were incremented in 5-dB steps, from ∼10 dB nHL up to 59.5 dB nHL. GSI Audera v2.7 software (Grayson-Stadler, Eden Prairie, MN, USA) was used to determine the ABR thresholds.

## Acknowledgements

We thank Dr. Yehoash Raphael, University of Michigan, for providing the *Gjb2^Pfl^* mice. We greatly appreciate molecular biology support and laboratory management by Bruce Derfler. We appreciate the use of the Nikon Ti2 inverted spinning disk confocal and Olympus VS200 Slide Scanner at the Harvard Medical School MicRoN Microscopy Core, and the Hitachi S-4700 scanning electron microscope at the Harvard Medical School Electron Microscopy Facility. This work was supported by a Blavatnik Biomedical Accelerator Pilot Grant, a Blavatnik Biomedical Accelerator Development Grant, a Q-FASTR Grant, and a Blavatnik Sensory Disorders Grant (from Harvard Medical School to D.P.C.); by a sponsored research agreement from WayVector, Inc. to D.P.C.; by a T32GM007748 Training Grant in Genetics fellowship to K.T.B.; by a Warren Alpert Fellowship to S.H.; and by a T32GM007753 MSTP Training Grant and Paul & Daisy Soros Fellowship for New Americans to M.A.N.

## Author contributions

M.V.I. conceptualization, data acquisition, data analysis, data interpretation, visualization of in vivo experiments in mice and NHPs, manuscript writing; K.T.B. conceptualization, data acquisition, data analysis, data interpretation, visualization of in vivo experiments in mice, manuscript writing; K.D.K. conceptualization, data analysis, data interpretation in mice; L.M.A. ABR data acquisition, data analysis, data interpretation; M.A.N., E.C.G., S.H., M.E.G. data acquisition, data analysis, data interpretation ATAC-seq experiments, AAV design, manuscript writing; C.W.P. data acquisition, data analysis, data interpretation of in vivo experiments in mice; S.P. data acquisition, AAV cloning; Y.L. data acquisition and neonatal injections in mice.; A.L. data acquisition, visualization of in vivo experiments in mice; D.P.C. conceptualization, supervision, data interpretation, manuscript writing.

